# Ribbon: Visualizing complex genome alignments and structural variation

**DOI:** 10.1101/082123

**Authors:** Maria Nattestad, Chen-Shan Chin, Michael C. Schatz

**Affiliations:** Cold Spring Harbor Laboratory, Cold Spring Harbor, NY, USA.; Pacific Biosciences, Menlo Park, California, USA.; Departments of Computer Science and Biology, Johns Hopkins University, Baltimore, MD, USA.

## Abstract

**To the Editor:** Visualization has played an extremely important role in the current genomic revolution to inspect and understand variants, expression patterns, evolutionary changes, and a number of other relationships^1–3^. However, most of the information in read-to-reference or genome-genome alignments is lost for structural variations in the one-dimensional views of most genome browsers showing only reference coordinates. Instead, structural variations captured by long reads or assembled contigs often need more context to understand, including alignments and other genomic information from multiple chromosomes.

---

We have addressed this problem by creating Ribbon (genomeribbon.com) an interactive online visualization tool that displays alignments along both reference and query sequences, along with any associated variant calls in the sample. This way Ribbon shows patterns in alignments of many reads across multiple chromosomes, while allowing detailed inspection of individual reads **(Supplementary Note 1)**. For example, here we show a gene fusion in the SK-BR-3 breast cancer cell line linking the genes CYTH1 and EIF3H. While it has been found in the transcriptome previously^4–6^, genome sequencing did not identify a direct chromosomal fusion between these two genes. After SMRT sequencing^7^, Ribbon shows that there are indeed long reads that span from one gene to the other, going through not one but two variants, for the first time showing the genomic link between these two genes (Figure 1a). More gene fusions of this cancer cell line are investigated in **Supplementary Note 2**. Figure 1b shows another complex event in this sample made simple in Ribbon: the translocation of a 4.4 kb sequence deleted from chr19 and inserted into chr16 (Figure 1b). Thus, Ribbon enables understanding of complex variants, and it may also help in the detection of sequencing and sample preparation issues, testing of aligners and variant-callers, and rapid curation of structural variant candidates **(Supplementary Note 3).**

**Figure 1.**
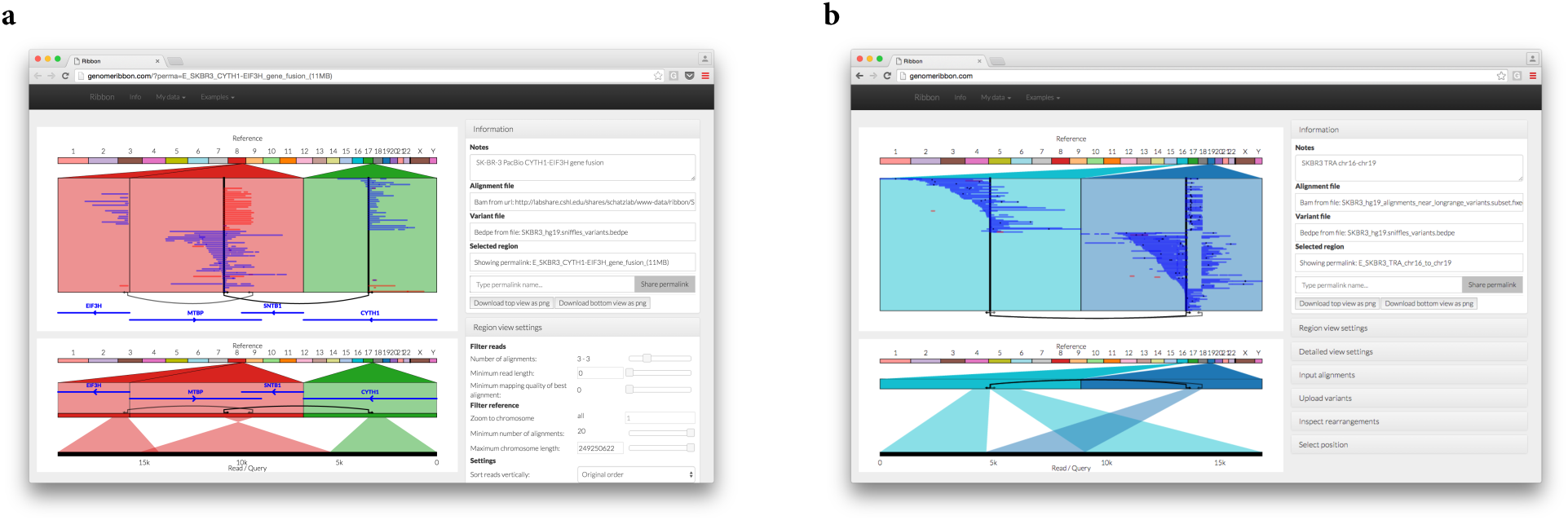
Ribbon shows all alignments from the perspective of both the reference and the read or query. Here long reads from SMRT sequencing show evidence for complex variants in the SK-BR-3 breast cancer genome. (a) The CYTH1-EIF3H gene fusion takes place through a series of two events that are captured within some of the long reads. (b) Long reads from SMRT sequencing show evidence of a full interchromosomal translocation where a sequence is homozygous deleted from chromosome 19 and homozygous inserted into chromosome 16.

In addition to SAM and BAM^8,9^ files with long, short, or paired-end reads, Ribbon can also load coordinate files from whole genome aligners such as MUMmer^10^. Therefore, Ribbon can be used to test assembly algorithms or inspect the similarity between species. **Supplementary Note 4** shows a comparison of gorilla^11^ and human genomes using Ribbon, highlighting major structural differences. In conclusion, Ribbon is a powerful interactive web tool for viewing complex genomic alignments.

## ACKNOWLEDGMENTS

This work was supported by NSF[DBI-1350041]; NHGRI[R01-HG006677]

## AUTHOR CONTRIBUTIONS

M.N. created Ribbon with input from all authors. M.N. and M.S. wrote the manuscript.

## COMPETING FINANCIAL INTERESTS

C.-S. C. is an employee and stockholder of Pacific Biosciences, a company commercializing DNA sequencing technologies. M.N. was a contractor of Pacific Biosciences.

